# Antibacterial efficacy of Jackfruit rag extract against clinically important pathogens and validation of its antimicrobial activity in *Shigella dysenteriae* infected *Drosophila melanogaster* infection model

**DOI:** 10.1101/2020.03.09.983015

**Authors:** NV Dhwani, Gayathri Raju, Sumi E Mathew, Gaurav Baranwal, Shivakumar B Shivaram, Neeraj Katiyar, Nilkamal Pramanik, Siddharth Jhunjhunwala, H.B. Shilpashree, Dinesh A. Nagegowda, Ritesh Kumar, Ajit K. Shasany, Raja Biswas, Sahadev A Shankarappa

**Affiliations:** Center for Nanosciences & Molecular Medicine, Amrita Institute of Medical Sciences & Research Center, Amrita University, Ponekkara P.O., Kochi 682041, India; Centre for BioSystems Science and Engineering, Indian Institute of Science, Bengaluru 560012, India; Molecular Plant Biology and Biotechnology Lab, CSIR-Central Institute of Medicinal and Aromatic Plants, Research Centre, Bengaluru - 560065, India; Biotechnology Division, CSIR-Central Institute of Medicinal and Aromatic Plants, P.O. CIMAP, Lucknow - 226015, India

**Keywords:** gut bacteria, antimicrobial, ingestion, antibiotic alternative, plant extract

## Abstract

The aim of this study was to determine the antibacterial property of extract derived from a part of the Jackfruit called ‘rag’, that is generally considered as fruit waste. Morpho-physical characterization of the Jackfruit rag extract (JFRE) was performed using gas-chromatography, where peaks indicative of furfural; pentanoic acid; and hexadecanoic acid were observed. *In vitro* biocompatibility of JFRE was performed using the MTT assay, which showed comparable cellular viability between extract-treated and untreated mouse fibroblast cells. Agar well disc diffusion assay exhibited JFRE induced zones of inhibition for a wide variety of laboratory and clinical strains of gram-positive and gram-negative bacteria. Analysis of electron microscope images of bacterial cells suggests that JFRE induces cell death by disintegration of the bacterial cell wall and precipitating intracytoplasmic clumping. The antibacterial activity of the JFREs was further validated *in vivo* using *Shigella dysenteriae* infected fly model, where JFRE pre-fed flies infected with *S. dysenteriae* had significantly reduced mortality compared to controls. JFRE demonstrates broad antibacterial property, both *in vitro* and *in vivo*, possibly by its activity on bacterial cell wall. This study highlights the importance of exploring alternative sources of antibacterial compounds, especially from plant-derived waste, that could provide economical and effective solutions to current challenges in antimicrobial therapy.

## Introduction

Antimicrobial agents including antiseptics and antibiotics are extensively used for infection control in community and nosocomial settings. Antiseptics such as chlorhexidine digluconate, triclosan and povidone-iodine are used as surface disinfectants in hand hygiene and disinfection of surgical and catheter insertion sites. In addition to human use, antibiotics such as ampicillin, amoxicillin, imipenem, and many others, are widely used to treat a variety of bacterial infections in animals, thus favoring their use in animal and poultry farming. Unfortunately, enhanced use of antimicrobial agents leads to the development of resistant microbes, such as chlorhexidine and colistin resistant *Klebsiella pneumonia*, methicillin and linezolid resistant *Staphylococcus aureus*, and vancomycin-resistant enterococci. Importantly, some of the antiseptics and antibiotics have also been reported to precipitate adverse systemic effects in patients^1^. Povidone-iodine and triclosan has been shown to disrupt thyroid hormone homeostasis^2^, while colistin and vancomycin have been associated with renal toxicity^3,4^. Thus, it is imperative that new and safe antimicrobial agents are explored from alternative sources. Plant extracts have been used for centuries to combat infectious human diseases in different parts of the world. Such medicinal extracts are a mixture of several compounds, with many extracts reported to have potent antimicrobial activities against wide range of drug resistant microbes^5^. Phenolics, terpenoids, alkaloids and lectins are some of the classes of compounds present in plant extracts that exhibit strong antimicrobial activity. Antimicrobial activity has been reported from various plant extracts such as *Brillantaisia lamium, Crinum purpurascens, Mangifera indica* and *Psidium guajava*, against variety of pathogens including *S. aureus, Enterococcus faecalis, Candida tropicalis, Cryptococcus neoformans*, and *Salmonella* Paratyphi. Thus, plant-based materials form an abundant source for antimicrobials, that could be economical, easy to process, and efficient against drug-resistant microbes. The Jackfruit is one such plant product that fits this description.

The fruit from the jack tree, botanically termed *Artocarpus heterophyllus*, belongs to the mulberry family, *Moraceae*. The fruit by itself is quite large, weighing approximately 4-10 kg, commercially inexpensive, and widely consumed in south-east Asia and Africa^6^. In addition to jackfruit’s nutritional value, various components of the fruit possess a plethora of medicinal properties^7^. The fruity arils are known to contain a cocktail of phytonutrients such as carotenoids^8^, isoflavones^9^, saponins^10^ and phenols^11^ that are responsible for antioxidative and immunomodulatory properties^7^. In addition, jackfruit leaf extract has been reported to possess antibacterial activities against *Escherichia coli, Listeria monocytogenes, Salmonella typhimurium, Salmonella enterica, Bacillus cereus, Enterococcus faecalis*, and *S. aureus*;^12^. Heartwood has antibacterial activities against *Bacillus subtilis, Streptococcus mutans, Streptococcus pyogenes, S. aureus* and *Staphylococcus epidermidis*^13^. In addition, extract derived from jackfruit seeds and shell powder have also been reported to demonstrate antibacterial activities against *L. monocytogenes*^14–16^.

While parts of the ripe jackfruit including the pulpy aril and seeds are a culinary delicacy, the rough, fibrous appendage called ‘rag’ that make up 10-20% of the fruit are either discarded as non-edible fruit waste, or in some cultures, cooked and consumed. There are no known reports of rag’s being used for medicinal purposes, although we have recently reported the use of the Jackfruit rag extract (JFRE) as a photo-sensitizer in solar cells^17^. Since rags make up large parts of the fruit and considering the existing lack of clarity towards their use for human benefit, we set out to explore possible medicinal value of the rag, with specific focus on its potential as an alternative to antibiotics against human pathogenic bacteria.

## Materials and methods

### Reagents

Methanol, 2-Propanol and hydrochloric acid (HCl) were purchased from Merck, India. 2, 2-diphenyl-1-picrylhydrazyl hydrate (DPPH), 3-(4, 5, dimethylthiazol2-yl)-2, 5-diphenyltetrazolium bromide (MTT), Glutaraldehyde and Triton-X 100 were purchased from Sigma Aldrich, USA. Dulbecco’s Modified Eagle’s medium (DMEM) and RPMI-1640 were purchased from Lonza, USA and fetal bovine serum (FBS) from Gibco, USA. Luria-Bertani (LB) and De Man, Rogosa and Sharpe (MRS) medium (broth and agar) were purchased from Himedia, India. Mouse dermal fibroblast (L929) cell line was obtained from National Center for Cell Science (NCCS), Pune. All experiments were performed in triplicate unless mentioned otherwise.

### Plant material and extract preparation

Rags were separated from jackfruit (*A. heterophyllus* Lam, obtained from a marked tree located in Shimoga district, Karnataka, India) and stored at −20°C until further use. Air dried rags were homogenized by grinding. Jackfruit rag extract (JFRE) was prepared by suspending the powdered rag (10% w/v) in 80% acidified (1.2 mol l^−1^ HCl) methanol and heated at 50°C for 5h, followed by addition of 100% methanol (1:2). Resulting solution was centrifuged (10,000 g, for 5 min at 4°C), and the supernatant was further concentrated using solvent evaporation techniques and lyophilized. The dried extract was then made up to a final concentration of 100 mg ml^−1^ in 100% methanol.

### Bacterial and fungal strains

All the bacterial strains including *Staphylococcus aureus* (SA113), *Mycobacterium smegmatis* (mc_2_155; ATCC700084), *Escherichia coli* (ATCC25922) and *Pseudomonas aeruginosa* (PA01; ATCC 15692), *Shigella dysenteriae*, *Klebsiella pneumoniae*, *Lactobacillus fermentum*, *Proteus vulgaris*, *Salmonella* Typhi and *Salmonella* Paratyphi A (Generously provided by Dr. Anil Kumar, Department of Microbiology, Amrita Institute of Medical Sciences and Research Center, Kochi, India) were cultured in LB broth at 37°C. *Candida albicans* (ATCC 2091) was cultured in Sabouraud dextrose (SD agar) at 37°C.

### Morpho-physical characterization of the jackfruit rag and its extract

Scanning electron microscopy: Surface morphology of rags was analyzed using a scanning electron microscope (JEOL, JSM-6490LA, Japan). Images were obtained after sputter coating samples with gold and imaging at an accelerating voltage of 15 kV. Elemental analysis of Jackfruit rag was performed using scanning electron microscope (SEM) equipped energy dispersive X-ray (ULTRA55/GEMINI, Zeiss, Germany), after drop casting the rag powder on silicon wafer.

UV-visible spectroscopy: Absorbance intensity of JFRE samples were measured from 400 to 700 nm wavelength using a uv-visible spectrophotometer (Synergy H1, Biotek, USA).

Fourier transform infrared (FTIR) spectroscopy: was performed on JFRE (Perkin Elmer spectrometer, L1860121, USA), by scanning samples from 4000-500 cm^−1^ for 32 consecutive scans at room temperature.

Raman spectroscopy: was performed on JFRE using LabRAM HR UV-VIS-NIR Raman microscope, Laser: 785 nm, Filter: D 0.6, Gratings: 600 IR, in the scanning range: 500-3000 cm^−1^.

Phytochemical tests on JFRE for detection of alkaloids, flavonoids, tannins, saponins and cardiac glycosides was performed according to previously published standard procedures^18^. (Supplementary, materials and methods)

### Gas chromatography–mass spectrometry

GC-MS quantification of methanolic RAG extract was carried out based on a previously published method^19^, using Agilent 7890 GC system equipped with auto-sampler, HP-5ms column and 7977A MSD mass detector (Agilent Technology). Briefly, 2 μl of diluted (1:100) sample (in pentane) was injected in splitless mode with Helium (He) as carrier gas. Inlet temperature was kept at 250°C and the column flow was set for 1.5 ml/min. The column oven was programmed for initial hold of 2 min at 40°C followed by 150°C with the ramp of 5 °C min^−1^. After 3 min hold at 150°C, the temperature was raised to 250°C with ramp of 5° C min^−1^ and 3 min hold at 250°C. Finally, the oven temperature was raised to 300°C with ramp of 10° C min^−1^and 5 min final hold at 300°C. MSD transfer line, MS Quad, and MS source temperature were kept at 280°C, 150°C and 230°C respectively. Spectra acquired in scan mode were processed analyzed and annotated using Mass hunter workstation software (Agilent Technology) and NIST 11 library.

### In vitro biocompatibility

#### Cell viability assay

L929 mouse fibroblasts were seeded in a 96-well plate at a density of 1×10^5^ cells per well and cultured under 5% CO_2_, at 37°C for 24 h in DMEM media supplemented with 10% FBS and 1% penicillin-streptomycin. After 24 h, the culture media was replaced with fresh media containing JFRE at concentrations of 0.1, 1, 10 and 100 mg ml^−1^. Viability of the JFRE-exposed L929 cells was determined using the MTT assay after 24 h. Solubilized formazan absorbance at 570 nm was measured using Biotek microplate reader (Synergy H1, Biotek, USA).

#### Hemolysis assay

Female Sprague-Dawley rats weighing around 200-250 g were euthanized using carbondioxide asphyxiation. Blood (5 ml) was collected in BD vacutainer tubes and further diluted with PBS (pH 7.4) in 1:10 ratio. Blood samples were centrifuged at 700 g for 5 min to separate erythrocytes from plasma, followed by addition of 0.1,1,10 and 100 mg ml^−1^ of JFRE in equal proportion. This mixture was incubated at 37°C for 1 h, followed by centrifugation at 700 g for 5 min. Peak absorbance values of the supernatant was measured at 540 nm (Synergy H1, Biotek, USA). Triton-X 100 (0.2%) treated erythrocyte suspension was used as a comparative positive control, and percent hemolysis in each sample groups was calculated relative to the positive control.

### Agar well disc diffusion assays

Antimicrobial activity of JFRE was evaluated using standardized agar well-diffusion assay^20^. Various microbial strains were plated, and 5 mm wells were created within agar plates. The wells were filled with either, JRFE (0.1, 1,10 and 100 mg ml^−1^), vehicle (80% methanol), or 1% penicillin-streptomycin solution (used as positive control for antibacterial activity assays) or Amphotericin-B (20 mg ml^−1^) used as positive control for antifungal activity assays), and incubated at 37°C for 24-48 h. Plates were imaged using Chemidoc^™^ Imager (Bio-Rad, USA), and zones of inhibition quantified using ImageJ software^21^. Minimum inhibitory concentration and minimum bactericidal concentration of JFRE against various bacterial strains were determined using the standardized broth dilution method as reported previously^22^.

### Fly experiments

Adult male *D. melanogaster* flies (4-5 d old) were raised in flasks containing classic banana-agar media, maintained at 60% humidity, and at 28°C. All feeding experiments were performed by placing JFRE (100 mg ml^−1^) or bacterial solution soaked (60 μl) circular filter paper discs on-top of banana-agar feed within each flask. *S. dysenteriae* cultured in 5 ml LB broth (16 h at 37°C), were pelleted and resuspended in PBS and subsequently used to infect flies. Four-hour pre-starved flies were allowed access to *S. dysenteriae*-infected feed for 24 h. Flies in the experimental group received JFRE feed, while the control groups received fresh banana agar. Flies were observed for 21 days and the number of surviving flies was counted each day.

### Statistical analysis

All data are shown as mean ± standard deviation (SD). Difference in mean values among various experimental groups were statistically tested using one-way ANOVA with post-hoc Bonferroni multiple comparison test, unless mentioned otherwise. A *‘p’* value of < 0.05 was considered as statistically significant.

## Results

### Physio-chemical characteristics of the jackfruit rag

On gross examination, the JFRs appear as yellowish-white fibrous bands with a smooth and shiny external surface, measuring approximately 5-6 cm in length, 0.5-0.7 cm in width, and weighing about 0.1-0.5 g (Fig. 1a). SEM images of an individual rag exhibited a ‘layered’ architecture (Fig. 1b), where dense ‘honey-comb’ patterned plant tissue (Fig. 1c) was interspersed between layers of fibrous sheets. The methanol extract itself was deep red (Fig. 1a, inset) in color and exhibited an absorbance maximum at 540 nm (Fig. 1d). Lyophilization of the extract yielded a dark-reddish powder. Using energy dispersive X-ray spectroscopy, it was determined that the rag extract powder comprised primarily of carbon and oxygen (Table S1), while other components such as Na, Cl and Si that were observed in the elemental analysis were most likely due to washing techniques and background signal from the silicone sample-loading wafer. Further chemical characterizations were performed using FTIR and Raman spectroscopy. In FTIR (Fig. 2a), the absorption peak at 3375 cm^−1^ likely corresponds to hydroxyl groups (-OH), and 1724 cm^−1^ indicates carbonyl (C=O) corresponding to a carboxyl or ester group arising from pectin or fatty acid residue^23^. The observed, peaks at cm^−1^ 1620 may be due to C=C stretch, and a group of stretching frequencies in the range of 1074 cm^−1^ – 1480 cm^−1^ potentially due to the presence of–CH_3_ and/or C–O group related to the flavonoids and hydroxyflavonoids^24^. In addition, peaks at 2926 cm^−1^ and 759 cm^−1^, could be due to the asymmetric stretching of an aliphatic (-CH_2_) group or bending mode of aromatic (=CH) group. Similarly, the 1450 – 1590 cm^−1^ bands observed in Raman spectroscopy (Fig. 2b) may be attributed to aliphatic bending vibration and/or aromatic stretching vibration. Additionally, the strong Raman band at 1235 cm^−1^ may be due to the twisting or rocking vibration of–CH_2_ group present in alicyclic or aliphatic compounds^25^.

**Fig. 1.**
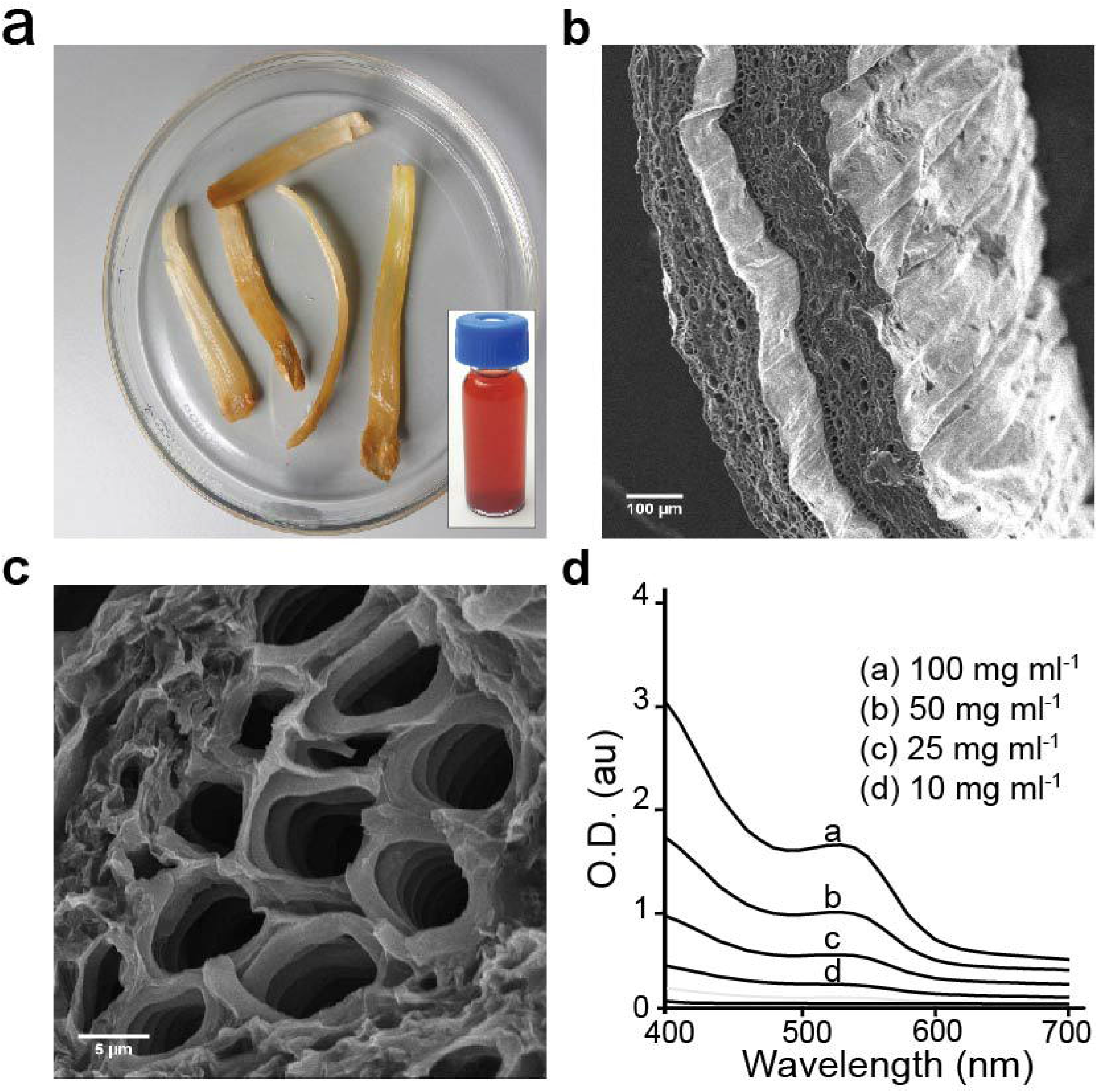
Characterization of jackfruit rag and extract. (a) Representative image of a peeled jackfruit exhibiting yellowish-white rags, and (inset) the deep-red methanolic rag extract. (b, c) Representative cross-sectional SEM images of a freshly harvested jackfruit rag. (d) UV-visible absorption spectra of varying dilutions of JFRE.

**Fig. 2.**
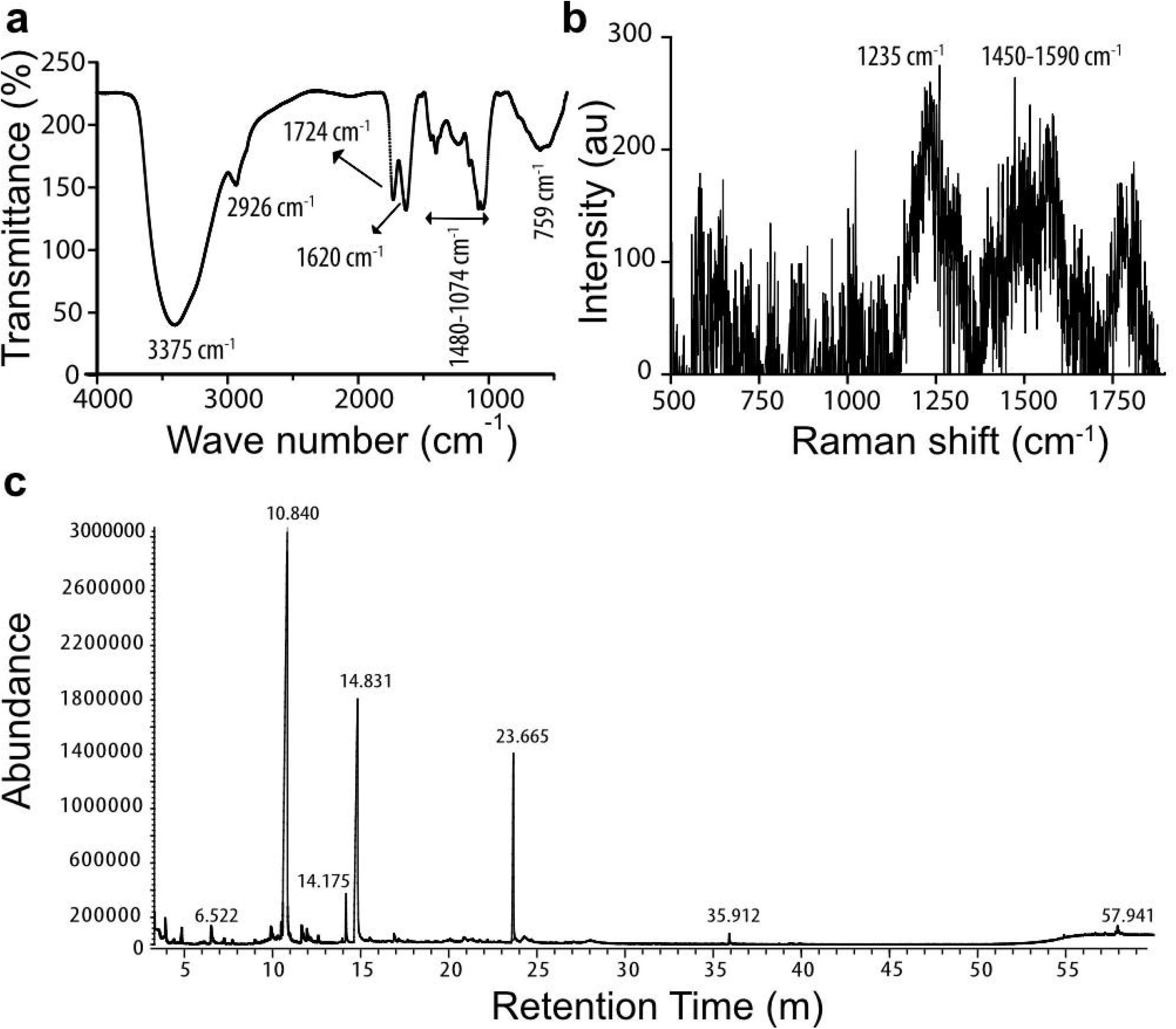
Spectroscopic analysis of jackfruit rag extract. (a) Fourier Transform Infrared (b) Raman spectroscopic analyses and (c) gas chromatography and mass spectroscopic studies to determine chemical composition of JFRE.

#### Gas chromatography mass spectrometry

To further identify the constituent compounds, the methanolic rag extract was subjected to GC-MS analysis. The spectral scan of JFRE annotated through Mass hunter workstation software (Agilent Technology) and NIST 11 library revealed six major constituent peaks which were identified as: furfural; pentanoic acid 4-oxo-methyl ester; levoglucosenone; 3-acetoxy-3-hydroxypropionic acid, methyl ester; citric acid, trimethyl ester; and hexadecanoic acid, methyl ester (Fig. 2c). The major contributing peaks calculated on the basis of peak area abundance were pentanoic acid, 4-oxo-, methyl ester (53.44%), 3-acetoxy-3-hydroxypropionic acid, methyl ester (27.06 %), and citric acid, trimethyl ester (8.91%) (Table 1).

**Table 1.** Major constituents present in methanolic RAG extract identified through GC-MS analysis

|  | RT | Compound name | Molecular Formula | Molecular Weight (g mol<sup>-1</sup>) | Peak area % |
| --- | --- | --- | --- | --- | --- |
| 1 | 6.52 | Furfural | C<sub>5</sub>H<sub>4</sub>O<sub>2</sub> | 96.021 | 1.33 |
| 2 | 10.84 | Pentanoic acid, 4-oxo-, methyl ester | C<sub>6</sub>H<sub>10</sub>O<sub>3</sub> | 130.063 | 53.44 |
| 3 | 14.17 | Levoglucosenone | C<sub>6</sub>H<sub>6</sub>O<sub>3</sub> | 126.032 | 2.20 |
| 4 | 14.82 | 3-Acetoxy-3-hydroxypropionic acid, methyl ester | C<sub>6</sub>H<sub>10</sub>O<sub>5</sub> | 162.053 | 27.06 |
| 5 | 23.66 | Citric acid, trimethyl ester | C<sub>9</sub>H<sub>14</sub>O<sub>7</sub> | 234.074 | 8.91 |
| 6 | 35.91 | Hexadecanoic acid, methyl ester | C<sub>17</sub>H<sub>34</sub>O<sub>2</sub> | 270.256 | 0.47 |

### Phytochemical assessment of JFRE

JFRE was subjected to an array of phytochemical screening tests (Supplementary, materials and methods) to determine the presence of common compounds that have been previously recognized to impart medicinal properties in plant extracts^26^. Results from phytochemical screening assays demonstrated the presence of polyphenolics, including anthocyanins, coumarins, and flavonoids. In addition, JFRE tested positive for compounds belonging to terpenoid, saponin and cardiac glycoside groups (Table 2).

**Table 2.**
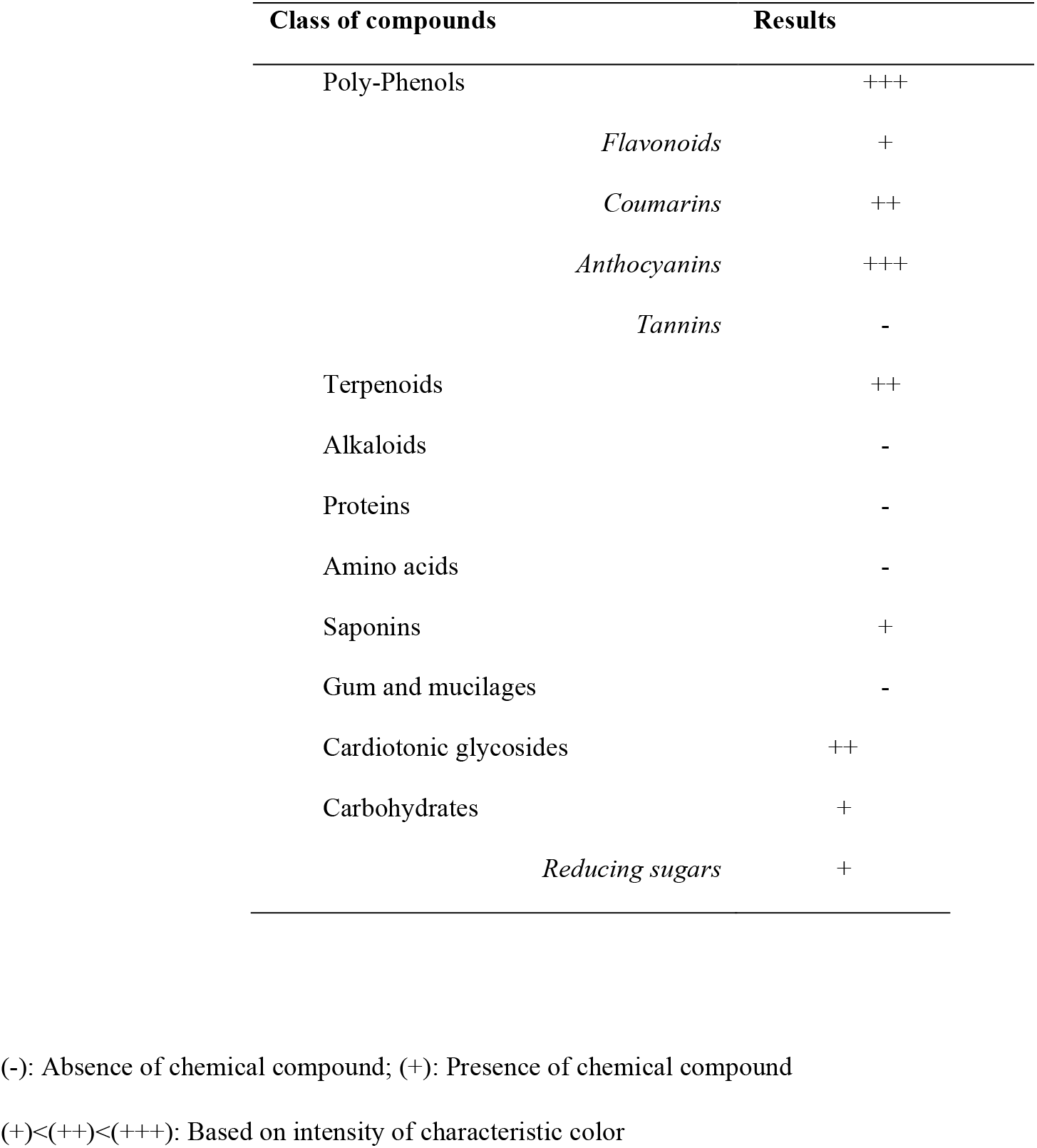
Biochemical analysis of JFRE.

#### In vitro biocompatibility testing of JFRE

To determine possible toxic effects, we examined the response of mouse fibroblast cell-line and rat erythrocytes to an acute exposure of JFRE. Viability of cultured fibroblast cells exposed to media containing 0.1,1, and 10 mg ml^−1^ of JFRE for 24 h was assessed using the MTT assay by measuring formazan absorbance at 570 nm. Interestingly, absorbance measurements from cultures exposed to JFRE were quite similar to those obtained from JFRE-free naïve cells (Fig. 3a), suggesting that the JFRE extract does not affect cellular viability in mammalian cells, at the concentrations tested. MTT measurements in cultures exposed to higher concentrations of JFRE (> 10 mg ml^−1^) resulted in unreliable absorbance signals, possibly due to extract induced interference in the assay, and hence higher JFRE concentrations were not tested.

**Fig. 3.**
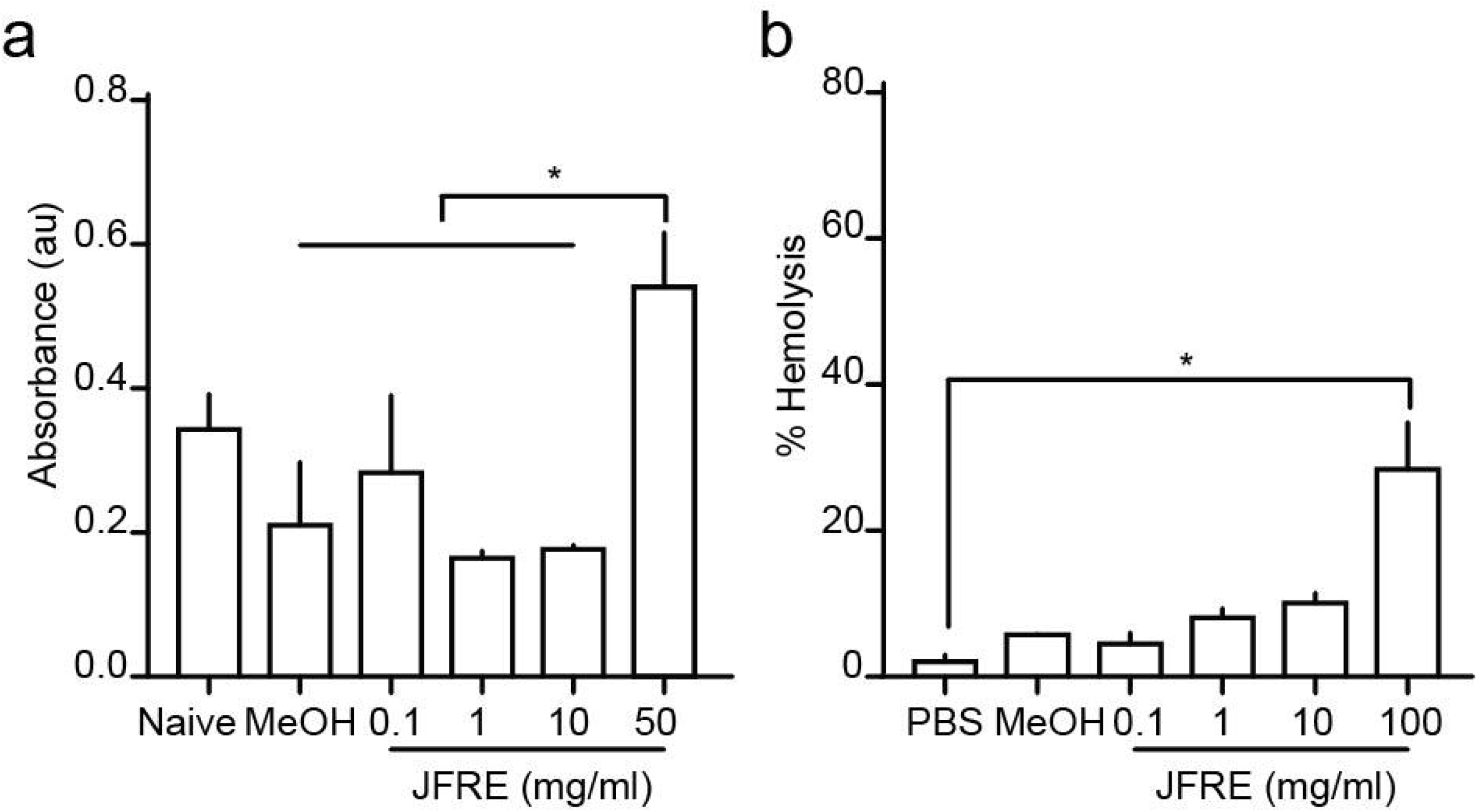
*In vitro* biocompatibility profile of the jackfruit rag extract. (a) Quantitative comparison of formazan absorbance values at 570 nm using the MTT assay from L929 cells exposed to varying concentration of JFRE for 24 h. (b) Quantitative comparison of hemolysis in erythrocyte suspension exposed to varying concentrations of JFRE. Percent hemolysis was calculated from peak absorbance values at 540 nm normalized to Triton-X induced hemolysis.

Similarly, blood collected from female Sprague-Dawley rats was processed to isolate erythrocytes, followed by the addition of 0.1, 1, 10 and 100 mg ml^−1^ of JFRE. Lysis of erythrocytes was assessed by measuring absorbance at 540 nm of the supernatant separated from the JFRE-erythrocyte mixture. JFRE-induced hemolysis was calculated by normalizing the resulting absorbance value against a completely hemolyzed erythrocyte control. We found that JFRE, at higher concentration induced 28% hemolysis (*p*<0.05, in comparison to all other groups), while lower concentrations produced hemolysis that was no different than PBS treated controls (Fig. 3b). These results suggest that JFRE possess hemolytic properties at higher concentrations, and hence caution needs to be exercised for potential applications related to intravenous use.

### Antibacterial property of JFRE

Antimicrobial activity of JFRE was assessed against a wide range of bacterial strains using agar well-diffusion and broth dilution assays. Interestingly, JFRE-exposed culture plates of both Gram-positive and Gram-negative bacteria demonstrated distinct zone of inhibition (Fig. 4a), suggesting broad-spectrum antibacterial activity of JFREs. Further, MIC and MBC values of JFRE against all bacteria tested were in the range of 5-10 and 10-20 mg ml^−1^ respectively (Table 3).

**Fig. 4.**
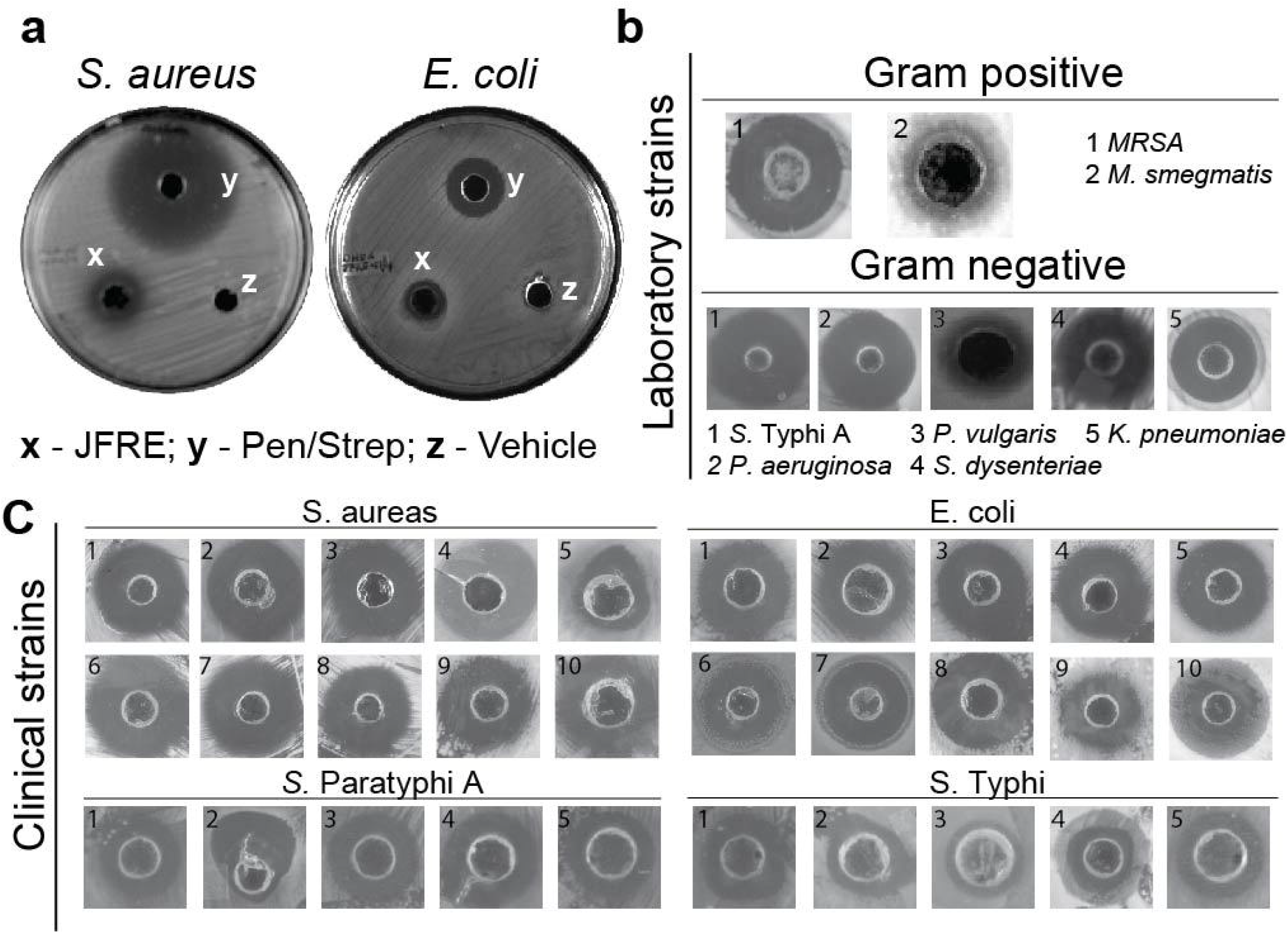
*In vitro* antibacterial activity of the jackfruit rag extract. (a) Representative images exhibiting clear bacterial growth inhibitory zones around agar wells containing JFRE (x) and an antibiotic (y) within well-plates that were cultured with laboratory strains of *S. aureus*, and *E. coli*. (b, c) Representative growth inhibitory zones from all laboratory and clinical bacterial strains tested in the study.

**Table 3.** Antimicrobial activity of JFRE against laboratory strains of bacteria.

| **Bacteria** | **MIC** | **MBC** |
| --- | --- | --- |
|  | (mg ml<sup>-1</sup>) | (mg ml<sup>-1</sup>) |
| <u>Gram positive</u> |  |  |
| *Staphylococcus aureas* | 5.0 | 10 |
| *Lactobacillus fermentum* | 7.5 | 15 |
| *Mycobacterium smegmatis* | 7.5 | 15 |
| <u>Gram negative</u> |  |  |
| *Escherichia coli* | 5.0 | 10 |
| *Shigella dysenteriae* | 5.0 | 10 |
| *Pseudomonas aeruginosa* | 5.0 | 10 |
| *Klebsiella pneumoniae* | 10.0 | 20 |
| *Proteus vulgaris* | 7.5 | 15 |
| *Salmonella Paratyphi A* | 7.5 | 15 |

Even though JFRE exhibited inhibition of laboratory bacterial strains (Fig. 4b), it is quite important that similar activity be examined in clinically relevant strains. Hence, we tested antibacterial activity of JFRE against strains of methicillin resistant *S. aureus*, *E. coli*, *S*. Paratyphi A and *S*. Typhi (Fig. 4c) derived from patient samples. Zones of inhibition similar to those observed in the laboratory strains were noticed. Since JFRE inhibited bacteria, irrespective of their strain, source, or antibiotic resistance, it is likely that the mechanism of action is different from conventional antibiotics^26,27^. Incidentally, JFRE did not appear to have antifungal effects against *C. albicans* (data not shown). Further, we imaged JFRE exposed cultures of *S. aureus* and *S. dysenteriae* using SEM. While vehicle treated bacteria exhibited symmetrical cells with regular borders and uniform morphology (Fig. 5a i, iii), JFRE treated bacteria appeared shrunken with irregular margins (Fig. 5a ii, iv). In addition, JFRE components appeared to have covered the bacterial cell surface, giving a matted appearance. Additionally, TEM images clearly showed the disintegration of bacterial surface, with formation of translucent intracellular clumped structures in JFRE treated *S. aureus* (Fig. 5b ii). These results suggest that JFRE could be producing its antimicrobial effects by direct activity on the outer bacterial cell wall or membrane.

**Fig. 5.**
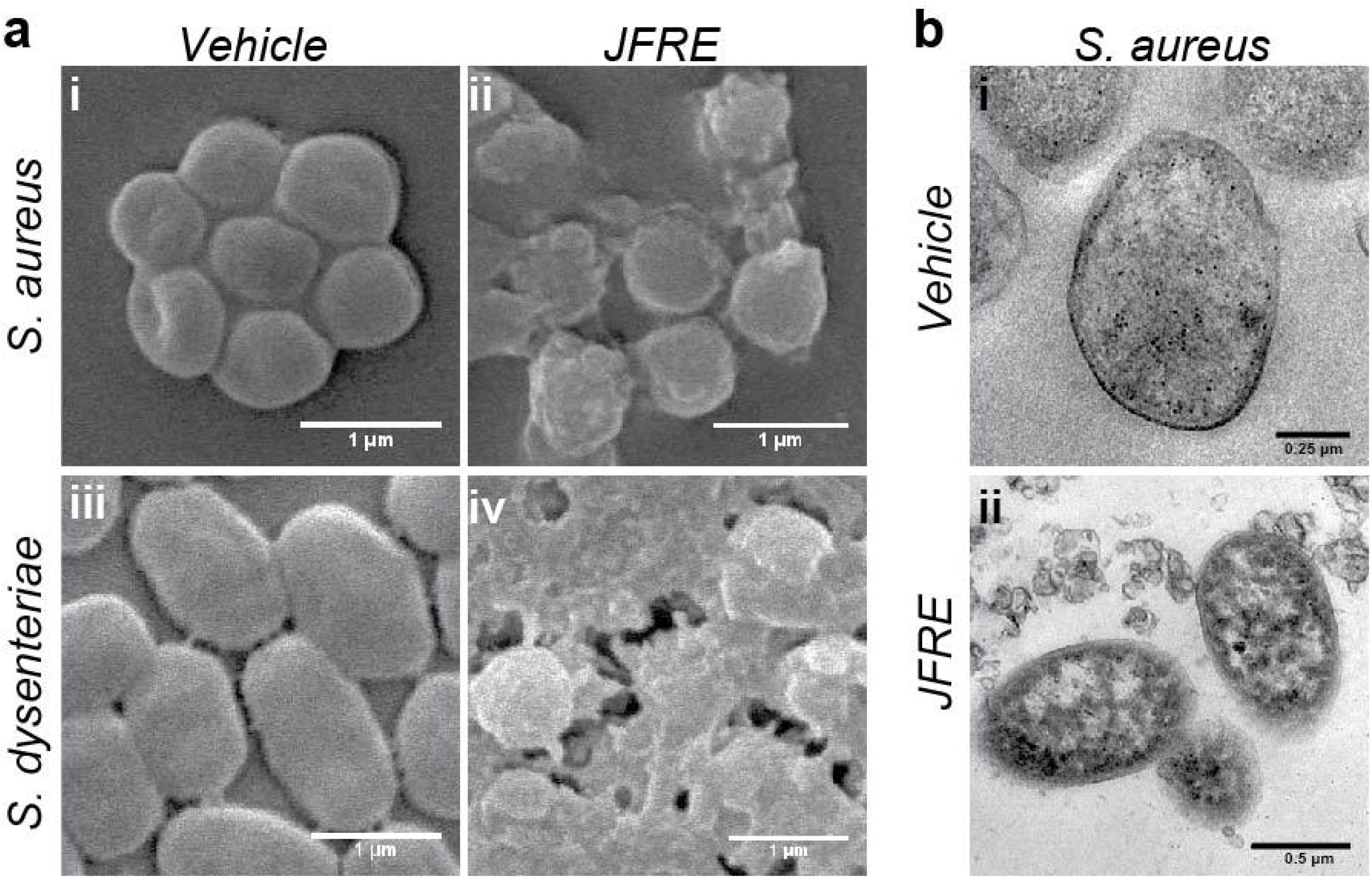
Electron microscope images of JFRE treated bacterial cells. Representative (a) SEM images of (i, iii) vehicle and (ii, iv) JFRE treated *S. aureus* and *S. dysenteriae*. JFRE treated bacteria exhibited irregular margins and had a ‘matted’ appearance. (b) TEM imaging of (ii) JFRE treated *S. aureus* showed loss of cell wall integrity (magnified segment from fig b, ii, inset) and dense cytoplasmic granularity, that was absent in (i) vehicle treated controls.

### Antibacterial effect of JFRE in *S. dysenteriae*-infected flies

To test the applicability of antibacterial property of JFRE in vivo, we challenged *S. dysenteriae* fed flies with JFRE and measured *D. melanogaster* survival rate for a period of 3 weeks. *S. dysenteriae*-fed flies exhibited a sharp drop (75%) in survival rate, starting 3-4 days after the infected feed (Fig. 6 iv). The survival rate continued to drop for the next 2 weeks and by the end of 15 days, the survival rate was 35%. No further reduction in survival rate was observed after two weeks. In stark contrast, *S. dysenteriae*-infected flies that were fed with the JFRE demonstrated a survival rate of 95% at the end of three weeks (Fig. 6 iii). These findings were quite similar to survival rates observed in uninfected flies that were treated with JFRE or vehicle alone (Fig. 6 i, ii), suggesting that JFRE protects against *S. dysenteriae* induced death in *D. melanogaster*.

**Fig. 6.**
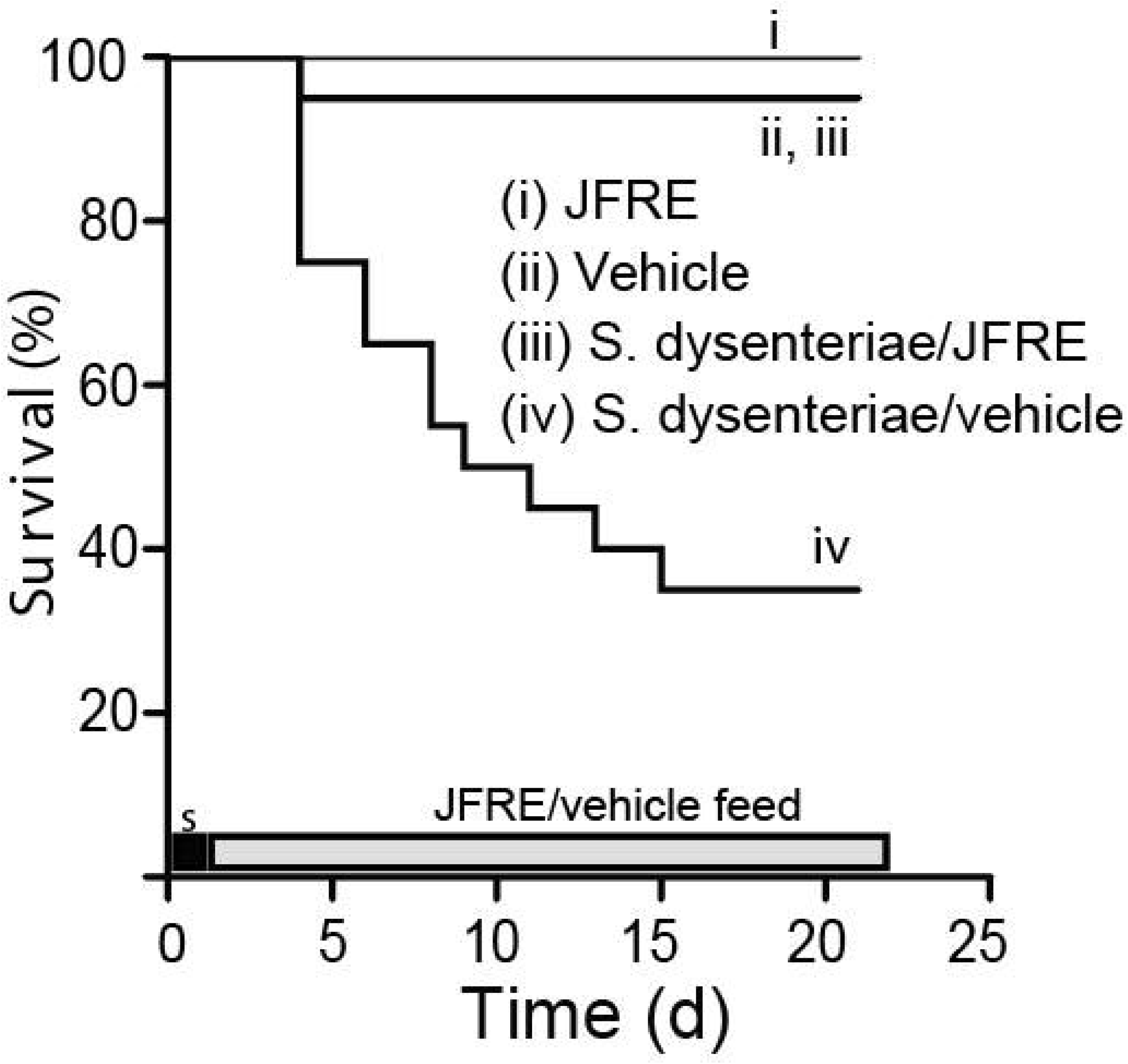
Survival of *S. dysenteriae* infected flies on JFRE treatment. Kaplan-Meier survival curves of *S. dysenteriae* pre-fed flies that were either fed JFRE, or vehicle alone. The dark bar on the x-axis (s) indicates the time duration of *S. dysenteriae* feed, while the lighter bar indicates the duration of JFRE/vehicle feed. n=20 per group. * p<0.05.

## Discussion

In this study, we set out to explore possible antimicrobial properties of JFREs. There have been several studies characterizing various components of the jackfruit^10^ and exhaustive lists have been compiled on their multiple uses^28^, to the best of our knowledge, we have not come across previous reports that have specifically focused on the jackfruit rag.

Morpho-physically, the yellowish-white rag has an interesting internal architecture, with each individual rag exhibiting an internal honey-combed pattern, along with a hydrophilic surface, considerable mechanical strength (tensile strength of 0.89 ± 0.4 MPa), and slow degradation in water (Figure S1). While the actual function of the jackfruit rag is not entirely clear, based on the structure and mechanical strength, it appears that it may be primarily used by the plant as a supportive, packing material to protect the fleshy aril and seeds. During the process of extract preparation, rag powder suspended in acidified methanol yielded deep red color, with an absorption maximum peak at 540 nm. This observation strongly suggested the presence of anthocyanins^29^, which was further confirmed using an array of biochemical screening tests. Further, the chemical characterization techniques of FTIR and Raman spectroscopy also suggested a mixture of carbon, oxygen and hydrogen containing compounds. Considering that the jackfruit belongs to the mulberry family which are well known for their high polyphenolic and anthocyanin content^30^, it would appear that jackfruit rag components also share similar characteristics. Incidentally, GC MS analysis revealed strong peaks indicating the presence of carboxylic acids, such as pentanoic acid and hydroxypropionic acid, that have been previously recognized to possess antibacterial properties^31–33^. Antioxidative property of JFRE was found to be moderate to low (data not shown) and was not further pursued.

We found that JFRE produced inhibitory zones in bacterial cultures at concentration of around 100 mg ml^−1^ (MIC 5-10 mg ml^−1^), while lower concentrations produced weak or no inhibitory zones in culture plates. While inhibitory zones were observed in both gram-positive and gram-negative laboratory strains, we were surprised to observe similar zones of inhibition in all the strains of clinically relevant cultures of *S. aureus, E. coli, S*. Typhi and *S*. Paratyphi A. This wide-ranging spectrum of antibacterial activity by JFRE against different bacterial strains, strongly points towards a possible target mechanism that could be common against all bacteria. SEM images clearly showed rag extract-induced blebbing of bacterial cells, along with aggregation, and change in shape, possibly due to membrane permeabilization. Previous studies have shown that bacterial cells that are normally impermeable to FITC, a fluorescent dye, become FITC-permeant^34^ after exposure to alcoholic extracts of various plant extracts mainly due to membrane permeabilization, destabilization, and disruption of membrane potential, resulting in cell blebbing and leakage of cellular contents^27^. In agreement with previous studies on other plant extracts^35^, our results also indicate that JFRE causes changes in bacterial cell morphology that eventually leads to cell death.

In view of these findings, we further tested the antimicrobial activity of JFREs in *S. dysenteriae* infected fly model. The protective effect of rag extract against *S. dysenteriae*-induced death was quite apparent, with 95% of the flies surviving for more than 3 weeks. This effect was consistently observed in all the three batches of fly cultures tested at different times. Considering that flies were pre-fed with *S. dysenteriae*-infected agar a day before rag-extract treatment, the rag extract appears to have directly prevented fly gut infection by eliminating *S. dysenteriae* and protecting flies from *S. dysenteriae* enterotoxin induced death. Further studies are however needed to identify the exact mechanism of JFRE-induced gut protection and its role in disrupting *S. dysenteriae* pathogenesis. It is also important to note that crude extracts, such as the one used in this study will contain a mixture of various bioactive molecules acting in cohesion to bring about a functional outcome, rather than a single active molecule exerting its effect^36^.

## Conclusion

In conclusion, we have demonstrated that the rag of jackfruit, which is generally considered as a non-edible fruit waste, has significant antibacterial activity. It is non-toxic when ingested and could possibly have applications in treating or preventing various drug resistant bacterial infections.

## Acknowledgments

We would like to acknowledge Dr. Anil Kumar, Department of Microbiology, AIMS, Kochi for generously providing bacterial strains. Department of Biotechnology, India for the Ramalingaswamy fellowship grant to Dr. Sahadev Shankarappa, and Department of Science and Technology for M. Tech grant to Ms. Dhwani N V. Dr. Nilkamal Pramanik is in receipt of the national postdoctoral fellowship and Dr. Siddharth Jhunjhunwala has received the Ramanujan Fellowship, both from the Department of Science and Technology. Govt. of India.

We would like to thank Mr. Kiran K.S for providing biochemical standards, Dr. Sabitha M for providing silica gel, Ms. Maneesha K. Suresh for data validation and technical help, Dr. Shantikumar V. Nair for critical comments on this manuscript, and Dr. Shivanee Shah for her valuable editorial support. Importantly, we would like to thank Mr. Adithya Shankar, Aramanekeri, for sourcing the raw materials for this study.

## Abbreviations

CFU: colony forming units
DPPH: diphenyl-1-picrylhydrazyl hydrate
FBS: fetal bovine serum
FTIR: Fourier-transform infrared spectroscopy
JFRE: jackfruit rag extract
MBC: minimum bactericidal concentration
MIC: minimum inhibitory concentration
MTT: 3-(4,5, dimethylthiazol2-yl)-2, 5-diphenyltetrazolium bromide
SEM: scanning electron microscope
GC-MS: Gas chromatography mass spectrometry.

---

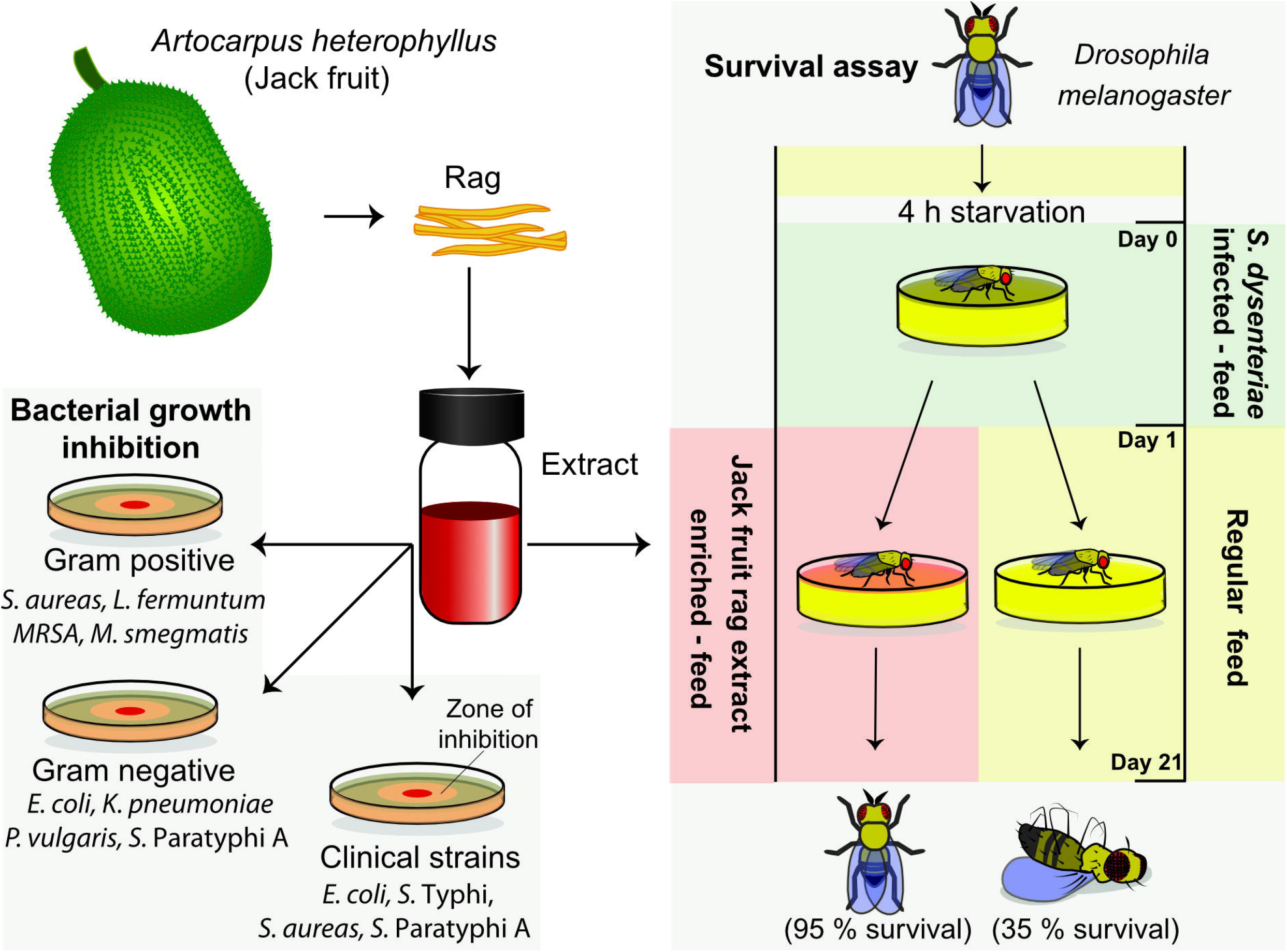

