## Supplemental material for "Antibacterial efficacy of Jackfruit rag extract against clinically important pathogens and validation of its antimicrobial activity in *Shigella dysenteriae* infected *Drosophila melanogaster* infection model"

### Supporting information

#### Materials and Methods

##### Phytochemical tests

An aqueous extract of JFRE was prepared by mixing concentrated JFRE in water at a concentration of 10 mg/ml. The suspension was vortexed, and centrifuged at 10,000 g for 10 min. The supernatant obtained (aqueous JFRE) was used for the following tests, unless mentioned otherwise.

*Test for amino acids – Ninhydrin test*

To test for amino acids, ninhydrin solution (Himedia, India) was added to aqueous JFRE in 1:25 ratio, and heated at 60° C for 15 min. Appearance of purple color was considered as an indicator for the presence of amino acids [1]. Bovine serum albumin (BSA) (Sigma, USA) was used as positive control.

*Test for Carbohydrate – Benedict's test*

Benedict's reagent was prepared by dissolving 173 mg $ml^{-1}$sodium citrate (Merck, India), 100 mg/ml sodium carbonate (Merck, India) and 17.3 mg $ml^{-1}$copper sulphate (Merck, India) in water. The prepared Benedict's reagent was added to aqueous JFRE in 1:1 ratio, and heated in a boiling water bath for 5 min. Appearance of a characteristic colored precipitate was considered as an indicator for carbohydrates [1]. Glucose (Merck, India) was used as a positive control.

*Test for reducing sugars – Fehling's test*

Filtered aqueous JFRE was added dropwise to Fehling's solution, and the mixture was boiled. Appearance of brick red colour was considered as indicator for presence of reducing sugars [2].

*Test for alkaloids – Mayer's test*

Mayer's reagent (Himedia) was added dropwise to concentrated JFRE and observed for formation of precipitate. Appearance of white precipitate in the solution was considered as indicator for presence of alkaloids [1].

*Test for polyphenols – Ferric chloride test*

Presence of polyphenols was tested by adding 50% v/v filtered aqueous JFRE to a mixture of 1:1 ferric chloride (1%) - potassium ferro cyanide (1%). Appearance of green and blue color was considered as an indicator for the presence of phenols and polyphenols respectively [2]. Pyrogallol was used as positive control.

*Test for tannins – Gelatin test*

Aqueous JFRE was mixed with 2% NaCl solution in 1:5 ratio, followed by dropwise addition of 1% gelatin. Formation of white precipitate was considered as an indicator of tannins [1,2].

*Test for flavonoids – Alkaline reagent test*

Ammonium hydroxide (10%) (Merck, India) was added dropwise to aqueous JFRE in a 1:5 ratio. Appearance of yellow color was considered as positive for flavonoids. Rutin was used as positive control.

*Test for coumarins*

Presence of coumarins in methanolic JFRE (10 mg $ml^{-1}$) was determined using thin layer chromatography (TLC) on silica gel using chloroform-methanol (9:1) as the solvent system [3]. Appearance of characteristic fluorescence under UV light (365 nm) was considered as positive for coumarins.

*Test for terpenoids – Salkowski test*

Aqueous JFRE, chloroform, and concentrated sulphuric acid were mixed in a ratio of 1:1:1.5 [4]. Appearance of reddish-brown color at the interface was considered as an indicator of terpenoids.

*Test for proteins – Biuret test*

Filtered aqueous JFRE was added to 2% copper sulphate in 20:1 ratio. Ethanol (95%) was added to the JFRE-$CuSO_4$ solution in 1:1 proportion, followed by addition of potassium hydroxide pellets until saturation was achieved. Appearance of pink color in the ethanolic layer was considered as an indicator for the presence of proteins.

*Test for cardiotonic glycosides*

Concentrated JFRE (25 mg $ml^{-1}$) was dissolved in chloroform, followed by addition of equal volume of concentrated $H_2SO_4$. Appearance of brown ring at the interphase was considered as an indicator for the presence of cardiotonic glycosides [2].

*Test for saponins*

Concentrated JFRE was dissolved in deionized water (2.5 mg $ml^{-1}$), and this mixture was shaken for 15 min. Formation of a layer of foam was considered positive for saponins [1].

*Test for gums and mucilages*

Aqueous JFRE was added to absolute alcohol in 1:2 ratio under stirring condition. Formation of white or cloudy precipitate was considered positive for gums and mucilages [1].

*Test for anthocyanins*

Dried JFR powder (100 mg $ml^{-1}$) was suspended in 80% methanol, followed by addition of 1.2 mol $l^{-1}$ HCl under 50°C. Formation of deep red color in the test solution was considered as a strong indicator for the presence of anthocyanins.

**Table S1.** The abundance of different elements (provided as atomic and weight percentages) in jackfruit rag extract as determined by the SEM-EDX spectral analysis. Experiment was performed once on different portions of a single sample and the mean values are provided.

| Element | Weight% | Atomic% |
|---|---|---|
| C K | 47.13 | 58.49 |
| O K | 33.48 | 31.19 |
| Na K | 8.03 | 5.21 |
| Si K | 3.04 | 1.61 |
| Cl K | 8.32 | 3.50 |
| Totals | 100 | 100 |

**Figure S1.** Graphical representation of change in weight of rag suspended in water for 30 days.

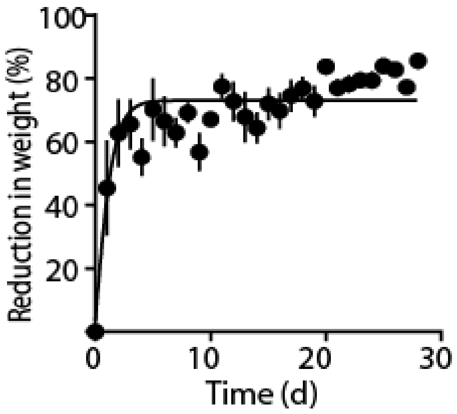
